# Suppression of Ovarian Cancer Cell Proliferation is Associated with Upregulation of Cell-Matrix Adhesion Programs and Integrin-β4-Induced Cell Protection from Cisplatin

**DOI:** 10.1101/2025.02.10.637535

**Authors:** Sadaf Farsinejad, Daniel Centeno, Jan-Savas Carstens, Teagan Polotaye, Tonja Pavlovic, Pouria Babvey, Taru Muranen, Pek Yee Lum, Laura A. Martin, Marcin P. Iwanicki

## Abstract

The role of extracellular matrix adhesion components in modulation of the treatment sensitivity of ovarian cancer (OC) cells is not well understood. Analysis of ovarian cancer TCGA gene expression data sets revealed an inverse correlation between genes involved in cell-cycle progression and extracellular matrix interactions including laminin-binding receptor integrin β4, a major component of extracellular matrix adhesion. Gene ontology analysis also showed that in patient populations with low integrin β4 expression, cell cycle-related programs were activated, while in populations with high expression of integrin β4, the activation of these cell cycle programs was lower. Suppression of proliferation with CDK4/6 inhibitor Palbociclib stimulated integrin β4 expression and induced protection against cisplatin in cells naturally expressing low levels of integrin β4. Additionally, ovarian cancer patient-derived organoids showed reduced cisplatin sensitivity when pretreated with Palbociclib. Our data also showed that integrin β4 overexpression decreased ovarian cancer cell proliferation and at the same time, attenuated cisplatin response. Our investigations reveal that expression of integrin β4 inversely correlates with cell cycle progression programs, whether observed in expression data of OC patient samples or in various OC cell lines. Consistently with these results, the overexpression of ITGB4 gene in ovarian cancer cell lines correlated with reduced cell proliferation rates and diminished sensitivity to cisplatin, supporting the idea that integrin β4 and likely its matrix ligands play critical roles in the regulation of cellular growth and chemoresistance of ovarian cancer cells.

## 1. Introduction

Recent characterization of the transcriptomic landscapes of ovarian cancer (OC) revealed an association of markers of enhanced proliferation with long-term survival^1^. These results suggested that the evolution of low-proliferating OC cell clones might be associated with the activation of growth-suppressing mechanisms that could restrict the actions of current therapies. We have recently published that chemotherapy-recovered ovarian tumors are associated with laminin deposition, and, in culture, elevation of predominantly laminin-containing extracellular matrix (ECM) deposition suppressed cell proliferation and the formation of OC outgrowths^2,3^. These studies raised an important question of whether ECM receptors (integrins) modulate the therapy response of slowly proliferating cancer cells.

Integrins are the cell surface αβ heterodimer glycoprotein molecules that bind ECM and transduce mechanical and biochemical signals from extracellular space into cells^4^. In vertebrates the combinatorial αβ complexity yields twenty-two distinct integrin dimers that can interact with various ECM components including collagens and laminins^5^. For instance, the Integrin β4 subunits can associate with the integrin α6 subunits to form laminin binding receptor^6^. Prior studies provide data that starvation of epithelial cells, which causes cell cycle arrest, induces integrin β4 expression and integrin β4/laminin-dependent survival^7^. Furthermore, recent experiments^8^ using triple negative breast cancer cell models have provided evidence that attenuation of integrin β4 expression induces DNA damage and evokes sensitization to Epirubicin, a DNA-damaging agent. While these studies established a possible link between integrin β4 expression, cell proliferation, DNA damage, and therapy response, the involvement of integrin β4 in supporting the evolution of growth-suppressed cancer cells and their response to therapy has not been previously explored in OC.

In this study, we used a combination of The Cancer Genome Atlas (TCGA) data analysis of ovarian serous cystadenocarcinoma, OC cell cultures, and RNA sequencing of growth-arrested OC cells to identify integrin β4 as a critical modulator of cell proliferation and cisplatin sensitivity. Our data are consistent with the model whereby laminin-binding receptors support the evolution of slowly proliferating, treatment-resistant ovarian cancer cell clones.

## 2. Materials and Methods

### Cell Culture

Our study used the following OC cell lines: HEYA8, KURAMOCHI, OVCAR4, OV90, CaOv3, TYK-nu, OVCAR3, OVSAHO, and RMUGS. KURAMOCHI, OVCAR4, OV90, and OVCAR3 cells were kindly provided by Dr. Denise Connolly (Fox Chase Cancer Center), HEYA8 cells was a kind gift from Dr. Sumegha Mitra (University of Indiana), TYK-nu, OVSAHO, and RMUGS cells by Dr. Joan Brugge (Harvard Medical School), and CaOv3 cells were purchased from the American Type Culture Collection (ATCC, Manassas, VA, USA). Each line was cultured under optimized conditions to support their growth and maintenance. KURAMOCHI, OVCAR4, OV90, and OVCAR3 cells were maintained in DMEM/F12 medium (Gibco, Waltham, MA, USA) containing 10% heat-inactivated fetal bovine serum (HI-FBS; Sigma-Aldrich, Burlington, MA, USA) and 1% penicillin-streptomycin (Gibco). HEYA8, TYK-nu, OVSAHO, RMUGS, and CaOv3 cells were cultured in a 1:1 mixture of MCDB 105 (Sigma-Aldrich) and Medium 199 (Gibco), supplemented with 5% HI-FBS and 1% penicillin-streptomycin. All cells were grown in a humidified 37 °C incubator with 5% CO_2_ and passaged at approximately 80% confluence using 0.25% trypsin-EDTA (Gibco). Mycoplasma testing was routinely performed every three to six months using the Uphoff and Drexler method^9^.

### 3D Spheroid Formation and Matrigel Supplementation

To generate 3D spheroids, 100 cells per well were seeded in a round-bottom, ultra-low attachment 96-well plate (Corning). After a quick centrifugation, the plate was incubated overnight so the cells could make their cell-cell adhesion and aggregate into spheroids. The next day, we allowed the plate to reach the room temperature. The Matrigel was thawed on ice at 4°C overnight and remained on ice to prevent premature gelation. We kept all tools and pipette tips cold (pre-cooled at −80°C). We added the Matrigel to a pre-cooled culture medium to make a concentration of 4% (v/v). Then this mixture was added to the spheroids making a final concentration of 2% (v/v). Subsequently, the spheroid plate was allowed to settle at room temperature before returning it to the cell culture incubator. This step ensured even distribution of Matrigel around spheroids before forming a gel. Spheroid cultures were maintained under normal culture condition for five to seven days after seeding, allowing for spheroid growth and potential outgrowth formation.

### Cell Proliferation Analysis

Cell proliferation in ovarian cancer cell lines was evaluated using the AlamarBlue assay (Invitrogen, #DAL1025) which measures metabolic activity of the cells and reflecting the cell proliferation. Each Cell line was seeded in 10 wells of a 96-well plate, with 5 wells for the initial time point (Time 1) after overnight attachment, and another 5 wells for 48 hours after the initial read (Time 2). The assay involves adding AlamarBlue reagent to each well, as per manufacturer’s instructions. The fluorescence was measured at excitation of 560 nm and emission of 590 nm using a SpectraMax i3x Microplate Reader (Molecular Devices, San Jose, CA, USA). The fluorescence values were background-subtracted, and results were normalized by dividing the fluorescence values at time 2 by those obtained at time 1 to calculate the relative proliferation rate.

Cell lines engineered for doxycycline-inducible expression of ITGB4, and their respective controls were seeded in black, ibidi-treated square plates to study the impact of Integrin β4 expression on cell proliferation. Both ITGB4-expressing and control groups were subjected to doxycycline for equal experimental conditions. Cell proliferation was monitored over a period of 48 hours using an automated fluorescent microscope, (BTLFX, Biotek, Lionheart, Winooski, VT, USA), taking pictures every 2 hours. For quantification, ImageJ (National Institutes of Health, Bethesda, MD, USA) software was used, combining automated cell detection with manual verification at the end to ensure accuracy. The proliferation rate was normalized to the initial number of cells so that growth dynamics could be directly compared under controlled conditions.

### Patient Derived Organoids

#### Organoid Derivation and Culture

Organoids were retrieved from the NYSCF ovarian cancer organoid biobank. Organoids were generated from fresh tumor tissue biospecimens obtained from patients diagnosed with ovarian cancer as described previously^10 11 12 13^. De-identified tumor material was provided by the NCI Cooperative Human Tissue Network (CHTN). Biospecimens correspond to remnants of tissue material that is removed as part of routine medical care collected in accordance with relevant state and local law. Other investigators may have received specimens from the same tissue specimens. The CHTN’s Research Resource identifier (RRID) is SCR_004446. For organoid derivation, tumor tissue was digested with 0.7 mg/mL collagenase (Sigma C9407) in the presence of 10 μM Y27632 (AbMole, cat. No. M1817) at 37°C for 25-50 minutes. Next, large undigested tissue fragments were removed by passing the tissue suspension through a 100 μm cell strainer. The filtered tissue was then centrifuged at 300g for 5 minutes. If any red blood cells were observed (red pellet) lysis was performed with red blood cell lysis buffer (Sigma-Aldrich, cat. no. 11814389001) for 3 minutes at room temperature and again centrifuged at 300g for 5 minutes. The dissociated tissue pellet was resuspended in ice-cold Cultrex Reduced Growth Factor BME type 2 (R&D Systems, cat. no. 3533-005-02) and at least 40,000 cells in 25 μL droplets were plated into pre-warmed 48-well plates. Droplets were allowed to solidify at 37°C for at least 30 minutes. Solidified droplets were then overlaid with ovarian tumor organoid medium containing 10 μM Y27632 (AbMole, cat. No. M1817). Organoid medium is composed of Advanced DMEM/F12 (Thermo-Fisher, cat. no. 12634-010), 1x GlutaMAX (Gibco, cat. no. 2492933), 1x Penicillin Streptomycin (10,000 U/ml) (Life Technologies, Cat. No. 15140122), 10 mM HEPES (Thermo Fisher, cat. no. 15-630-080), 100 mg /mL Primocin® (InvivoGen, cat. no. ANT-PM-1), 1x B-27™ supplement (Gibco, cat. no. 17504-044), 1.25 mM N-Acetyl-L-cysteine (Sigma-Aldrich, cat. no. A9165), 10 mM Nicotinamide (Sigma-Aldrich, cat. no. N0636), 0.5 mM A83-01 (Tocris, cat. no. 2939), 0.5 mg /mL Hydrocortisone (Sigma-Aldrich, cat. no.H0888), 10 mM Forskolin (R&D systems, cat. no. 1099), 100 nM β-Estradiol (Sigma-Aldrich, cat. no. E2758), 16.3 mg/mL Bovine Pituitary Extract (Thermo Fisher, cat. no. 13028014), 10 ng/mL recombinant human FGF-10 (PeproTech, cat. no. 100-26), 5 ng/mL recombinant human FGF-7 (PeproTech, cat. no. 100-26), 37.5 ng/mL recombinant human Heregulin Beta-1 (PeproTech, cat. no. 100-03), 5 ng/mL recombinant human EGF (PeproTech, cat. no. AF-100-15), 0.5 nM WNT Surrogate-Fc fusion protein (ImmunoPrecise, cat. no. N001), 100 ng/mL R-Spondin1 (PeproTech, cat. No. 120-38) and 1% Noggin-Fc fusion protein conditioned medium (ImmunoPrecise, cat. no. N002). Media was changed every 2-3 days and organoids passaged every 7-10 days.

#### Drug Assays in Patient Derived Organoids

Grown organoids ranging 30-90 μm in diameter were dissociated with TrypLE Select (Thermo Scientific, cat. No. 12563-011) containing 10 μM Y27632 and 10 μg/mL of DNAse I (Sigma, cat. no. DN25) for 10 minutes at 37°C. TrypLE solution was then neutralized with Advanced DMEM/F12 containing 1x Gibco™GlutaMAX (ThermoFisher Scientific, 35050-079), 1x Penicillin Streptomycin (10,000 U/ml) (Life Technologies, Cat. No. 15140122), 10 mM HEPES (ThermoFisher Scientific, 15-630-080), 100 μg/mL Primocin® (InvivoGen, ant-pm-1), 10 μM Y27632 and 10 μg/mL DNAse I. Next, to aid dissociation into single cells, organoids were mechanically sheared using a 1 mL syringe with 30-gauge needle. Dissociated organoids were centrifuged at 300g for 5 minutes at 4°C. Pellets were then resuspended at 100 cells/μl in ice-cold 70% Cultrex® Reduced Growth Factor BME, Type 2, PathClear™ (Cultrex BME) (R&D Systems, Cat. No. 3533-010-02) diluted in organoid medium containing 10 μM Y27632 and 10 μg/mL DNAse I, and kept on ice until use. Inside a tissue culture hood, the single cell solution was then plated into cell-repellent, clear-bottom 384-well plates (Greiner Bio-One, cat. no. 781976) using an *Assist Plus* pipetting robot (Integra, cat. no. 4505) with *Voyager II* automatic pipette (Integra, cat. no. 4732). During dispensing, source plates were kept on an integrated active cooling block (Inheco, cat. no. 7000190). Using a pre-programmed plating protocol (VIALAB software), 10 μL of cell suspension was dispensed to each assay well for a final density of 1000 cells per well. Assay plates were then placed into an incubator for 10 minutes at 37°C to allow the Cultrex BME to polymerize. After 10 minutes, plates were returned to the tissue culture hood and 20 ml of organoid media containing 10 mM Y27632 and 10 mg/mL DNAse I was added to each assay well with Integra pipetting robot, as above. After 4 days, 20 ml of organoid media without Y27632 or DNAse I was added to each assay well using Integra pipetting robot as above. After feeding, 100 nM or 400 nM of Palbociclib (Selleckchem, cat. no. S1116) was added to treatment wells and 0.1% DMSO (Sigma, cat. no. D2650) to vehicle control wells of the assay plates using an I.DOT S automated liquid handler (Dispendix). 72 hours later, using the I.DOT liquid handler, Cisplatin (Selleckchem, cat. no. S1166) was added in a 9-point serial dilution with doses ranging between 22.9 nM to 150 μM to both Palbociclib-treated and vehicle control wells. In parallel, additional wells were treated with 10 μM Staurosporine (Selleckchem, cat. no. S1421) as a killing control. Next, the plates were sealed with a semi-porous membrane (Sigma-Aldrich, cat. no. Z380059) and placed in the incubator at 37°C, undisturbed for 5 days. At the end of drug treatments, cell viability was quantified using CellTiter-Glo 3D Cell Viability Assay (Promega, cat. no. G9683) following manufacturer’s directions. Luminescence values were quantified using a Clariostar Plus plate reader (BMG Labtech, cat. no. 0430-501).

#### Data analysis and statistics

IC_50_, R^2^ and AUC values for each drug treatment were calculated with GraphPad Prism (version 10.0.3). Results are presented as means ± SD. The number of independent biological or technical replicates is indicated. The statistical significance was determined by ordinary One-way ANOVA and Tukey’s multiple comparisons test with a single pooled variance, as indicated in each figure.

### Bioinformatics Analysis of TCGA Ovarian Cancer Transcriptomic Data

#### Data Retrieval and Preprocessing

RNA sequencing data was downloaded from The Cancer Genome Atlas (TCGA), specifically for the ovarian cancer project (TCGA-OV). This data was accessed using the TCGAbiolinks package, part of the Bioconductor software (version 3.14), an open-source project that provides tools and packages for the genomic analysis in the R programming environment^14^. This data included STAR Counts which is a gene expression quantifications method. The raw data was formatted into an expression matrix using the GDCprepare function. To ensure comparability across samples, the expression data were normalized to Transcripts Per Million (TPM) using the DESeq2 package which adjust for variations in sequencing depth and gene length^15^. Using the tidyverse package, expression data from individual patients were combined into single dataset. This dataset included TPM values, aligned gene identifiers and patient identifiers, and the combined data was then saved in a TSV format for subsequent analyses.

#### t-SNE Visualization of Transcriptomic Data

Utilizing the Rtsne package in R, the t-Distributed Stochastic Neighbor Embedding (t-SNE) was performed. This method was applied to visualize patterns of gene expression across ovarian cancer samples. In the t-SNE plot predefined gene sets associated with GOBP_CELL_MATRIX_ADHESION^16^ and KEGG_CELL_CYCLE^17^, were highlighted with colors, depicting a non-quantitate perspective of transcriptomic data structure and the relationships among these two gene sets.

#### Gene Set Variation Analysis (GSVA) and Correlation Analysis

Gene Set Variation Analysis was executed using the GSVA package in R. This method delivers a non-parametric, unsupervised method to estimate gene set enrichment across the samples^18^. First, the two gene sets, GOBP_CELL_MATRIX_ADHESION” and “KEGG_CELL_CYCLE” were extracted from the Molecular Signatures Database (MSigDB)^19 20^. Then the enrichment scores were derived for each gene set across the samples. Having the GSVA scores, pairwise correlations between gene sets were calculated using Pearson’s correlation method. A heatmap was then generated to visually represent these correlations using ggplot2 package. finally, hierarchical clustering was applied to cluster similar gene sets on the heatmap.

#### Analysis of Cell Cycle Gene Expression by ITGB4 Levels in Ovarian Cancer

To explore the differential expression of cell cycle genes in regard to ITGB4 expression in ovarian cancer, we stratified samples into three groups based on the quantiles of ITGB4 expression levels. Annotating the top 20th percentile for high expression, the 20th to 80th percentiles for mid expression, and the lowest 20th percentile for low expression. Following the sample classification, a heatmap was generated to visually represent the expression profiles of cell cycle-related genes across the high, mid, and low ITGB4 expression groups.

#### Gene Ontology (GO) Enrichment Analysis

Subsequently, a Gene Ontology (GO) enrichment analysis was done to determine the biological processes that were highly upregulated or downregulated in the high ITGB4 group versus low ITGB4 group. This analysis used the clusterProfiler package and identified significant GO terms with p-values adjusted for multiple testing using the Benjamini-Hochberg method^21^. Finally, the significant GO terms were visually represented in bar plots using the enrichGO function for each of the two categories of upregulated and downregulated gene sets.

#### Heatmap of RNA-seq Differential Gene Expression (DEG)

To visually analyze the differential expression of DEGs in palbociclib-treated spheroids compared to controls, we utilized the pheatmap package in R. The gene expression data were first preprocessed to ensure only genes with significant differential expression were included. The expression data were then normalized to minimize variance; this normalization involved scaling each gene’s expression levels by subtracting the mean and dividing by the standard deviation of that gene across samples. A heatmap was generated to display the normalized data. Clustering was applied to the rows (genes) to group genes with similar expression patterns.

To focus on specific biological processes across the RNA-seq data, gene sets related to “GOBP_CELL_MATRIX_ADHESION” and “KEGG_CELL_CYCLE” were retrieved from the Molecular Signatures Database (MSigDB). The entire gene expression matrix was filtered to include only the selected gene set, after which the heatmap was generated as described above.

The code for the analyses described above is available on GitHub at https://github.com/pbabvey/cellProliferation_matrixAdhesion.

### RNA-Seq of Palbociclib-Treated Spheroids

#### Cell Treatment and Spheroid Formation

HEYA8 cells were pre-treated with 500 nM palbociclib (Selleckchem, cat. no. S1116) or the carrier control for 72 hours. Following treatment, cells were seeded into ultra-low attachment plates as spheroids in a growth medium containing 2% Matrigel, as previously described in this paper methodology for 3D Spheroids and Matrigel Supplementation. Spheroids were allowed to grow for seven days under standard culture conditions.

#### RNA Extraction and Quality Assessment

Total RNA was extracted from spheroids using the E.Z.N.A.® Total RNA Kit (Omega Bio-tek, Norcross, GA, USA) according to the manufacturer’s protocol. The concentration and purity of RNA were evaluated using a NanoDrop 2000 spectrophotometer (Thermo Fisher Scientific). The RNA samples were subsequently stored at −80 °C freezer before being shipped on dry ice to Genewiz (Azenta US, South Plainfield, NJ, USA) for RNA sequencing.

#### RNA Sequencing Workflow

The RNA sequencing was sent to GENEWIZ (Azenta Life Sciences). The process started with PolyA-based mRNA enrichment. Then mRNA fragmentation and cDNA synthesis were performed through first and second strand priming. Subsequent steps included end-repair, 5′ phosphorylation, and adenine (dA) tailing followed by adaptor ligation. The libraries were enriched using PCR. Sequencing was performed on an Illumina HiSeq 2500 device, producing paired-end 150 bp reads. The raw sequencing data underwent quality assessment and trimming to remove adapters and low-quality bases using Trimmomatic v0.36. Next, the cleaned reads were aligned to the Homo sapiens GRCh38 reference genome (ENSEMBL) using STAR v2.5.2b. Gene hit counts were obtained using featureCounts from the Subread package v1.5.2. The resulting count data was used for subsequent differential expression analyses.

#### Differential Expression Analysis

Using DESeq2, a comparison of gene expression between palbociclib-treated and control samples was performed. Statistical significance was determined through the Wald test, which computed p-values and log2-fold changes. Genes with adjusted p-value (Padj) < 0.05 and an absolute log2-fold change > 1 were called as differentially expressed genes (DEGs).

#### Gene Set Enrichment Analysis

Gene set enrichment analysis (GSEA) was performed using GSEA_4.2.2 software^22^ against selected gene sets from the Molecular Signatures Database (MSigDB 7.5)^19^. The Signal2Noise metric was applied for ranking, and 1,000 permutations were used. Results with a false discovery rate (FDR) < 0.25 were considered significant.

#### Plasmids and Cloning

First, the pCW57.1 vector (Addgene plasmid, #41393) which contains a TRE (Tetracycline Response Element) promoter allows for the inducible expression of genes in the presence of tetracycline or its analog doxycycline. The pCW57.1 vector was modified by inserting a T2A peptide fused to a fluorescent protein marker, (mAG monomeric Azami-Green). The T2A-mAG cassette was PCR amplified from the pBOB-EF1-FastFUCCI-Puro plasmid (Addgene plasmid, #12337). Primers were designed to introduce overhangs compatible with NEBuilder HiFi DNA Assembly (New England Biolabs, Ipswich, MA, USA). The PCR product and the pCW57.1 vector were assembled using Gibson Assembly to generate the intermediate vector, Assembled_pCW_T2A. The assembled PCR products were then purified from the agarose gel using the Qiagen Gel Extraction (Kit Qiagen, Hilden, Germany). The Integrin β4 coding sequence was PCR amplified from the pRc/CMV-Beta(4)Integrin (Addgene plasmid, #16265) using a primer pair designed to introduce overhangs complementary to the T2A sequence present in the Assembled_pCW_T2A vector. Resulting PCR products and the Assembled_pCW_T2A backbone were co-assembled using NEBuilder HiFi DNA Assembly to ultimately yield the construct PCW_T2AMAG_ITGB4_Assembled. This construct included a bicistronic design with mAG which is reporter gene downstream of the Integrin β4 sequence, separated by a T2A peptide to allow for their concurrent expression. The expression of GFP fluorescence in these cells served as an indicator of successful Integrin β4 overexpression.

#### Plasmid Transformation and Expansion

The constructed plasmid is transformed to competent E. coli cells by the method of heat shock for bacterial transformation. Transformation colonies were selected on LB agar plates containing ampicillin. The positive colonies were grown in LB broth and then plasmids isolated with a standard miniprep kit (Qiagen, Hilden, Germany). To check the correctness of the assembled construct, enzymatic digestion was and thereafter electrophoresis was performed. The resulting fragments were analyzed on an agarose gel to confirm correct assembly based on size and the pattern of expected fragments.

#### Lentivirus Production

Lentivirus was prepared by transfection of HEK293T with mixture of packaging plasmids psPAX2 (Addgene #12260) and pMD2.G (Addgene, #12259) with the lentiviral vector PCW_T2A-MAG-ITGB4_Assembled and incubated in serum-free Opti-MEM (Gibco) and Lipofectamine 3000 (Invitrogen). The medium was refreshed the following day. The viral particles were collected from the supernatant of the cell culture at 48 and 72 hours post-transfection. The viral supernatant was then collected and used for the transduction of target ovarian cancer cell lines, HEYA8, and OV-90. Transduction was performed by adding viral supernatant and polybrene (Santa Cruz Biotechnology, Dallas, TX, USA) to cell culture at a final concentration of 10 μg/mL. Following transduction, the cells were incubated in normal growth conditions for 24 to 48 hours before selection pressure was applied by using an appropriate antibiotic. Post-selection, the presence and expression of Integrin β4-T2A-MAG fusion protein were confirmed by Western blot analysis with and without doxycycline (Sigma-Aldrich, #D9891) induction.

#### Western Blot Analysis

Cells were washed twice with PBS, lysed on ice-cold RIPA buffer (Cell Signaling Technology), supplemented with Halt™ Protease Inhibitor Cocktail (Thermo Fisher), then collected using a cell scraper while kept on ice. Cell lysates were pelleted at 20,000× g for 20 min, the supernatant was collected, and the protein concentration was determined by BCA assay (Pierce) according to the manufacturer’s instructions. The lysates were then combined with 6X sample buffer, boiled at 95°C for 10 min, followed by loading and resolution by electrophoresis on a polyacrylamide gel. Proteins were then transferred to Immobilon® PVDF transfer membranes (Millipore Sigma, Burlington, MA, USA), which were blocked with 5% non-fat milk in Tris-buffered saline containing 0.1% v/v Tween-20 (TBST) for 30 minutes at room temperature. Afterwards, membranes were incubated with primary antibodies overnight at 4 °C in TBST containing 5% non-fat milk. Membranes were then washed three times with TBST and incubated in an HRP-conjugated secondary antibody (1:10,000) for 1 h at room temperature. Membranes were then washed three times in TBST. Membranes were developed using ImmobilonTM Forte enhanced chemiluminescent substrate (Millipore Sigma, Burlington, MA, USA) and visualized using an iBright CL1500 (Thermo Fisher). The following primary antibodies were used in this study: Integrin β4 (LSBio, #LS-C357939), GAPDH (Santa Cruz Biotechnology, #sc-47724). HRP-conjugated secondary antibodies were purchased from Santa Cruz Biotechnology (Dallas, TX, USA).

#### Flow Cytometry

We performed flow cytometry to measure the ITGB4 expression on the cell surface of OC cells. Cells were harvested by trypsinization, collected by centrifugation, and washed twice with PBS containing 2% FBS. An automated cell counter was used to count the cells (5x10^5 to 1×10^6) and stained with monoclonal ITGB4 antibody (LifeSpan BioSciences, # C357939) at 0.75 μg per 10^6 cells and incubated for 1 hour at room temperature. Cells were then washed twice with PBS, fixed in 4% paraformaldehyde (PFA) for 15 minutes and washed again. Cells were incubated with Alexa Fluor 568 secondary antibody (Invitrogen, #A-11011) at a 1:500 dilution and followed by washing and resuspending in PBS. In parallel, cells stained only with the secondary antibody were used as control to determine the background signal. To determine cell viability adherent cells were trypsinized, centrifuged at 300× g for 5 minutes, and washed twice with PBS containing 2% FBS. Cells were then incubated on ice with PBS containing 2% FBS and 2 μg/mL propidium iodide (PI) (Molecular Probes, Eugene, OR, USA). Data acquisition was performed using an Attune NxT Flow Cytometer (Thermo Fisher) and analyzed with FlowJo software (Version 10.10.0, FlowJo, Ashland, CA, USA). The cell death was reported as the percentage of PI-positive cells and cell viability was reported as percentage of PI-negative cells.

#### Statistical Analysis

Statistical analyses were done using GraphPad Prism 9.0 (GraphPad Software, San Diego, CA, USA) and statistical significance was determined with unpaired, two-tailed, parametric t-tests and adjustment for multiple comparisons (Benjamini-Hochberg method), one-way ANOVA followed by Tukey’s multiple comparisons test, or a two-way ANOVA. In all the statistical analysis p ≤ 0.05 was considered statistically significant.

## 3. Results

### 3.1. Examinations of ovarian cancer TCGA data sets reveal an inverse correlation between cell-matrix adhesion and cell cycle gene expression

To determine the relationship between cell cycle regulation and cell-extracellular matrix (ECM) adhesion in OC, we analyzed 433 TCGA ovarian cancer transcriptomes downloaded from the Genomic Data Commons (GDC) Data Portal through custom R scripts. These transcriptomes represent the most prevalent subtype of ovarian cancer, ovarian serous cystadenocarcinoma, which primarily includes high-grade serous ovarian carcinoma^23^. We employed the t-distributed Stochastic Neighbor Embedding (t-SNE) method to visualize the expression landscape of genes involved in both the cell cycle and cell-matrix adhesion pathways. This dimensionality reduction technique helped us reveal clusters that represent distinct expression patterns of these gene groups across the patient cohort. By overlaying specific gene markers from the Molecular Signature Database (MSigDB; https://www.gsea-msigdb.org/gsea/msigdb/), specifically the gene sets GOBP_CELL_MATRIX_ADHESION and KEGG_CELL_CYCLE on the t-SNE plot, we observed clear clustering patterns, indicating the potential relationship between these gene sets (**Fig. 1A**). To further elucidate the relationship between the cell cycle and cell-matrix adhesion genes, we conducted a correlation analysis, calculating the pairwise correlations between these two sets of genes. Using the Gene Set Variation Analysis (GSVA), we identified a negative correlation between the expression of cell-matrix adhesion genes and many of the genes regulating cell cycle progression (**Fig. 1B**). Notably, our pairwise analysis demonstrated a negative correlation between cell cycle genes and laminin-binding integrin β4 (**Fig. 1B** arrow). Interestingly, our previously published experiments provided evidence that increasing deposition of laminin-rich ECM suppresses cell proliferation^2^. These results suggest the possibility that integrin β4 expression levels vary depending on the cell cycle status in ovarian cancer cells. In addition to the predominant negative correlation with integrin β4, several genes positively correlated with cell cycle regulators (red portion in **Fig. 1B**). Some of these genes, such as PTPRC (CD45), LILRB1, and IKZF1 are involved in immune response, suggesting a potential role in immune signaling in cell proliferation^24 25^. Genes such as FAP and INHBA are linked to ECM remodeling and fibrosis, which may support proliferation^26^. These findings highlight the interplay between immune signaling, the microenvironment, and cell cycle progression.

**Figure 1.**
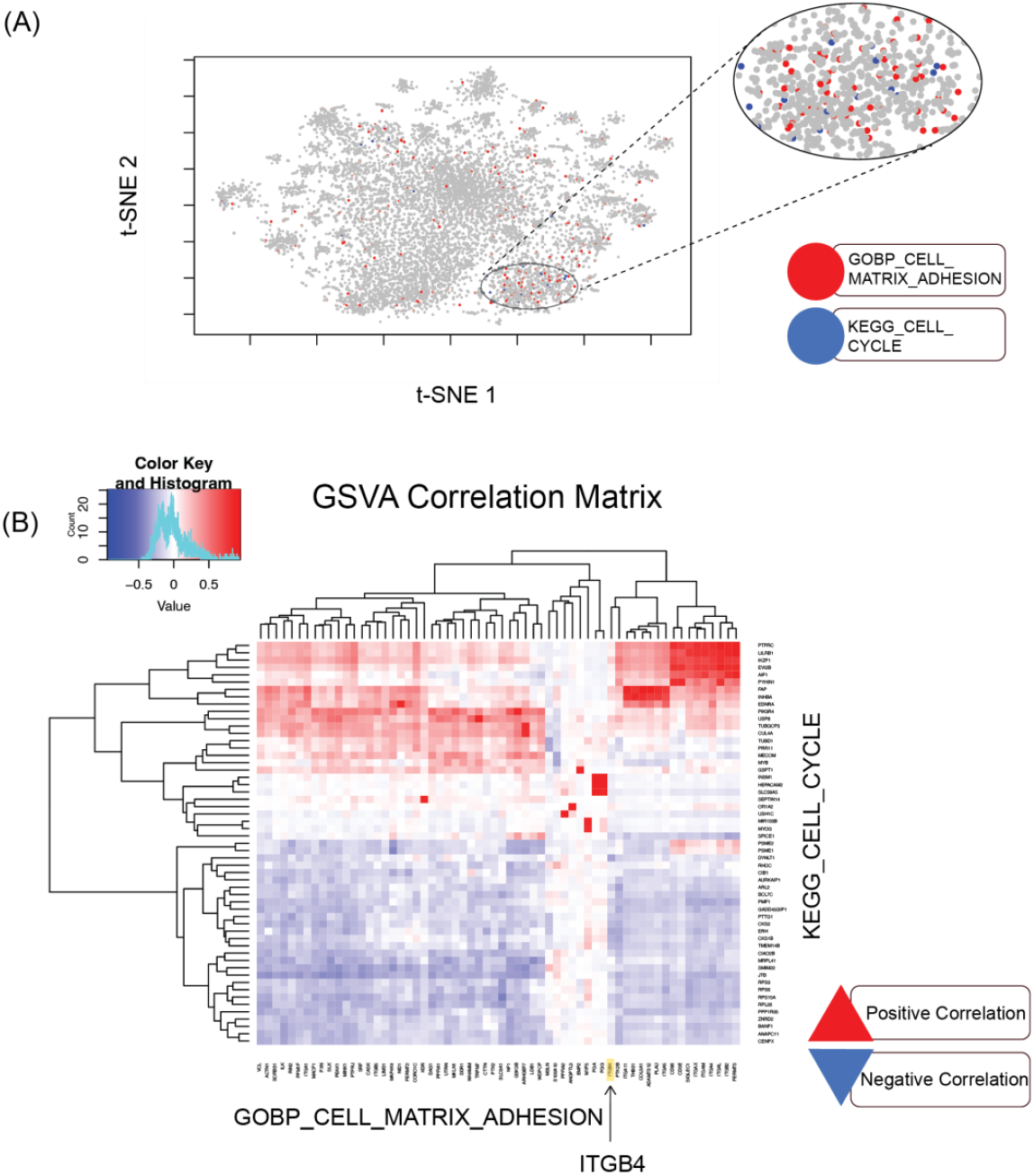
Inverse correlation between cell-matrix adhesion and cell cycle gene expression in high-grade serous ovarian carcinoma. (**A**) t-SNE visualization of variance-expressed genes across 433 patient transcriptomes from TCGA-OV. The magnified oval highlights colocalization of cell-matrix adhesion and cell cycle genes, suggesting their connection. (**B**) Correlation analysis between cell cycle and cell-matrix adhesion genes using Gene Set Variation Analysis (GSVA). This analysis demonstrates a significant negative correlation between cell cycle genes and cell-ECM adhesion molecules, including integrin β4 (arrow).

### 3.2. Integrin β4 mRNA expression changes with cell cycle genes in ovarian cancer patients

To test the possibility of whether integrin β4 mRNA expression levels change in relation to the ovarian cancer cell cycle, we analyzed gene expression profiles across The TCGA ovarian cancer datasets. Patients were categorized based on their ITGB4 mRNA expression into high, medium, and low groups. Our heatmap analysis (**Fig. 2A**) reveals a striking inverse correlation where high ITGB4 expression is associated with the downregulation of some cell cycle genes, suggesting a potential suppressive role of ITGB4 on cell cycle progression. Conversely, lower ITGB4 expression levels are linked with upregulated cell cycle gene expression, indicating a more active cell cycle.

**Figure 2.**
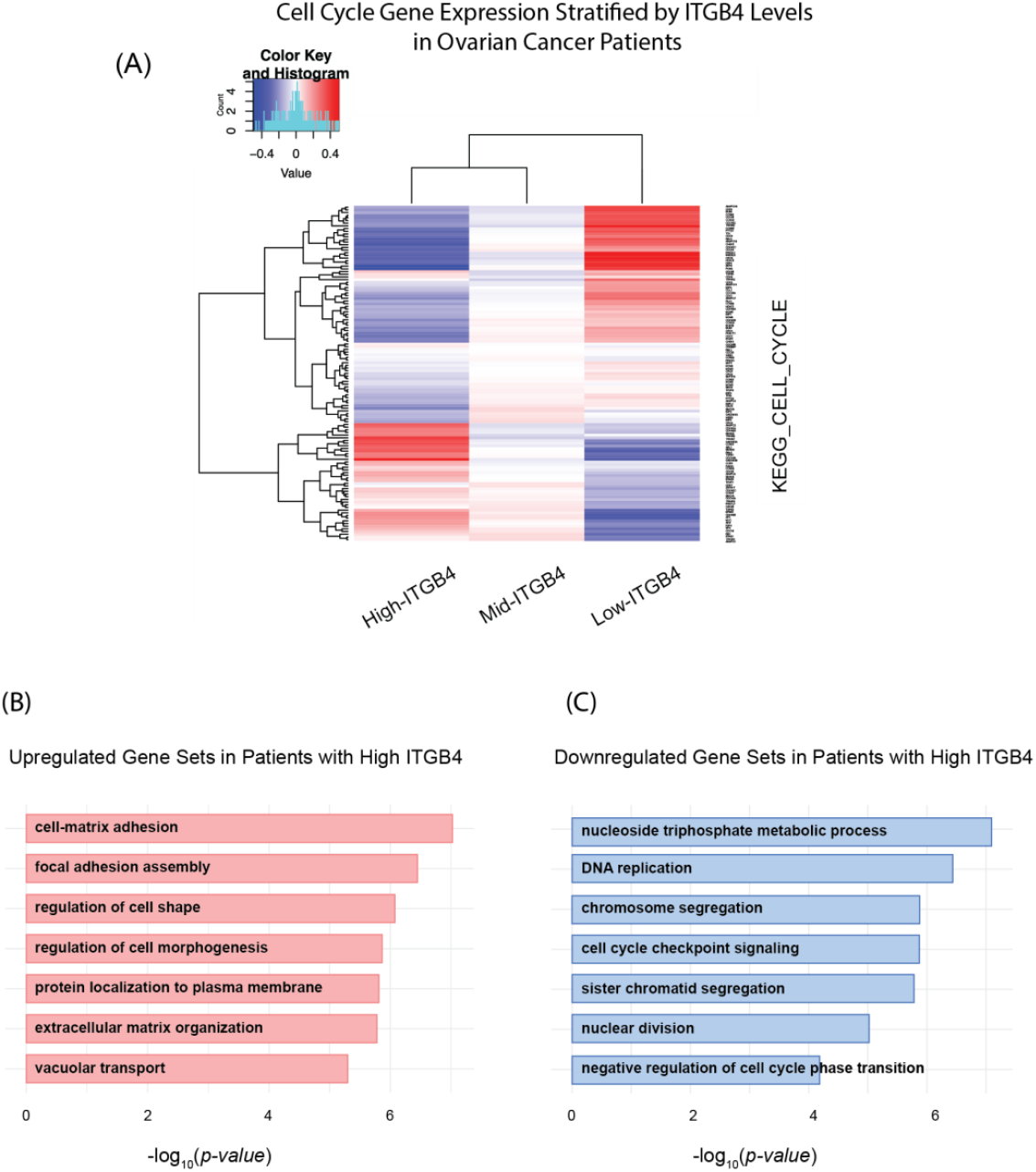
Differential expression and functional impact of ITGB4 in Ovarian Cancer. (**A**) Heatmap of cell cycle gene expression across ovarian cancer patients, categorized by ITGB4 mRNA levels. High ITGB4 (top 20 percentiles) is associated with generally lower expression of cell cycle genes (KEGG_CELL_CYCLE), whereas low ITGB4 (bottom 20 percentiles) is linked to broadly higher expression of these genes. (**B**) Upregulated gene sets in patients with high ITGB4. Statistical analysis via t-tests and adjustment for multiple comparisons (Benjamini-Hochberg method) identified cellular processes such as cell-matrix adhesion and extracellular matrix organization that are significantly upregulated. (**C**) Downregulated gene sets in patients with high ITGB4. Similar statistical methods reveal significant downregulation in processes related to cell cycle progression and DNA replication, indicating a potential shift towards a more quiescent cellular state.

Furthermore, our gene ontology analysis of ovarian tumor samples with high integrin β4 expression highlighted several critical cellular mechanisms being influenced. Specifically, we observed a pronounced activation of gene sets associated with cell-matrix adhesion and extracellular matrix organization, suggesting a pivotal role for integrin β4 in modulating the interaction between ovarian cancer cells and their microenvironment (**Fig. 2B**). Conversely, the analysis also revealed a significant downregulation in gene sets related to core cell cycle processes, such as DNA replication, chromosome segregation, and cell cycle checkpoint signaling (**Fig. 2C**). This pattern suggests that high integrin β4 levels may contribute to a reduction in cell proliferation rates, possibly through inducing a more quiescent cellular state. These findings raised the question of whether high integrin β4 protein expression is inversely correlated with OC cell proliferation.

### 3.3. Integrin β4 protein expression inversely correlates with OC cell proliferation

To determine whether integrin β4 protein expression inversely correlates with cell proliferation, we probed integrin β4 protein expression, and measured cell proliferation rates among various OC cell lines including OVCAR4, OV90, KURAMOCHI, CaOv3, TYK-nu, RMUGS, HEYA8, OVSAHO, and OVCAR3. Western blot analysis revealed the following integrin β4 expression levels in ovarian cancer cell lines: lowest in HEYA8, highest in RMUGS and CAOV3, and intermediate in OVCAR4, OV90, KURAMOCHI, OVSAHO, OVCAR3 and Tyk-nu (**Fig. 3A**). Our observations also revealed an inverse relationship between integrin β4 expression and cell proliferation: cells with lower integrin β4 levels (HEYA8) exhibited the highest proliferation rates, while those with higher levels (RMUGS) showed the least (**Fig. 3B**). These findings support the hypothesis that integrin β4 expression is associated with the regulation of cell proliferation.

**Figure 3.**
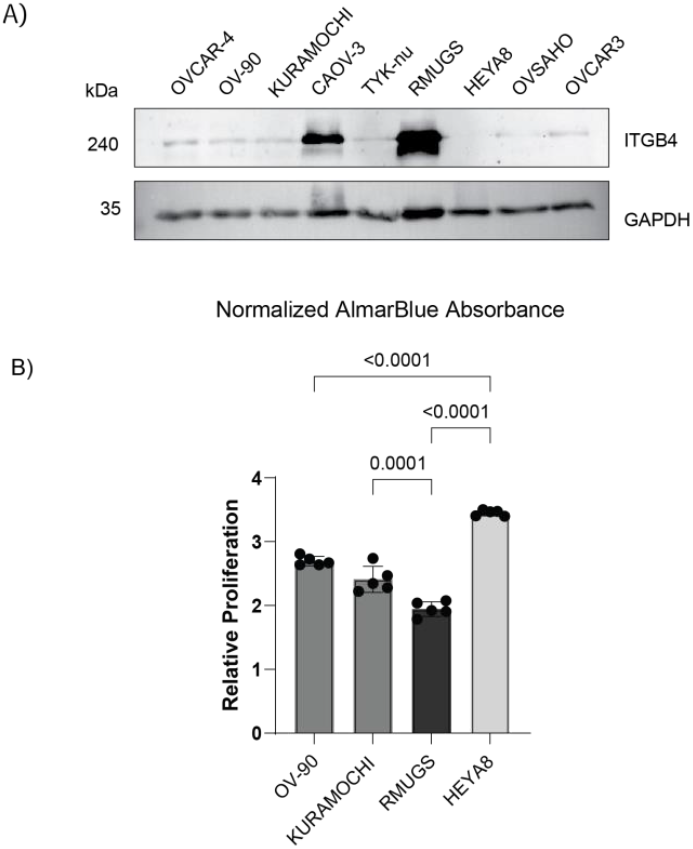
Correlation between ITGB4 protein expression and cell proliferation in ovarian cancer cell lines. (**A**) Western blot analysis of integrin β4 protein in various OC cell lines, including OVCAR4, OV90, Kuramochi, CaOv3, TYK-nu, RMUGS, and HeyA8. (**B**) Proliferation rates measured by normalized AlamarBlue fluorescence across cell lines, showing inverse correlation with ITGB4 expression. Each dot represents an independent experimental replicate. Statistical analysis was performed using one-way ANOVA followed by Tukey’s multiple comparisons test, indicating significant differences in proliferation rates between cell lines with different levels of ITGB4 expression (p < 0.0001).

### 3.4. Inhibition of OC cell proliferation with Palbociclib activates cell-ECM adhesion programs, expression of integrin β4 and induces protection from cisplatin

So far, our data provide evidence that laminin receptor integrin β4 expression correlates with low OC cell proliferation. Next, we wanted to explore the possibility of whether directly inhibiting cell cycle progression, in cells expressing low levels of integrin β4, stimulates ECM adhesion programs and integrin β4 expression. To examine this possibility, we treated HEYA8 (Integrin β4^low^) with Palbociclib (Ibrance), an FDA-approved cyclin-dependent kinase 4/6 (CDK4/6) inhibitor. We also observed that in three-dimensional ECM-supplemented suspended HEYA8 cultures, Palbociclib significantly decreased the formation of HEYA8 outgrowths, which are multicellular extensions protruding from spheroids (**Fig 4A**). Taken together these data indicate that Palbociclib-treated HEYA8 cells downregulate cell cycle progression programs and inhibit outgrowth formation.^2^ Next, we wanted to know whether inhibition of cell proliferation with Palbociclib promotes the expression of cell matrix-adhesion genes. To do this we performed RNA sequencing of suspended HEYA8 cell cultures reconstituted with 2% Matrigel and treated with vehicle control or 500 nM Palbociclib (**Fig. 4A**). RNA sequencing revealed that over 2,600 genes were differentially expressed (DEGs) in Palbociclib-treated HEYA8 spheroids compared to controls which highlights the broad impact of Palbociclib on gene regulation (**Fig. 4B**). As expected, Palbociclib treatment significantly downregulated the expression of genes associated with cell cycle progression (**Fig. 4C**). Interestingly, we observed that Palbociclib induced the expression of cell-matrix adhesion and cell-matrix receptor genes, including integrin β4 (**Fig. 4D**).

**Figure 4.**
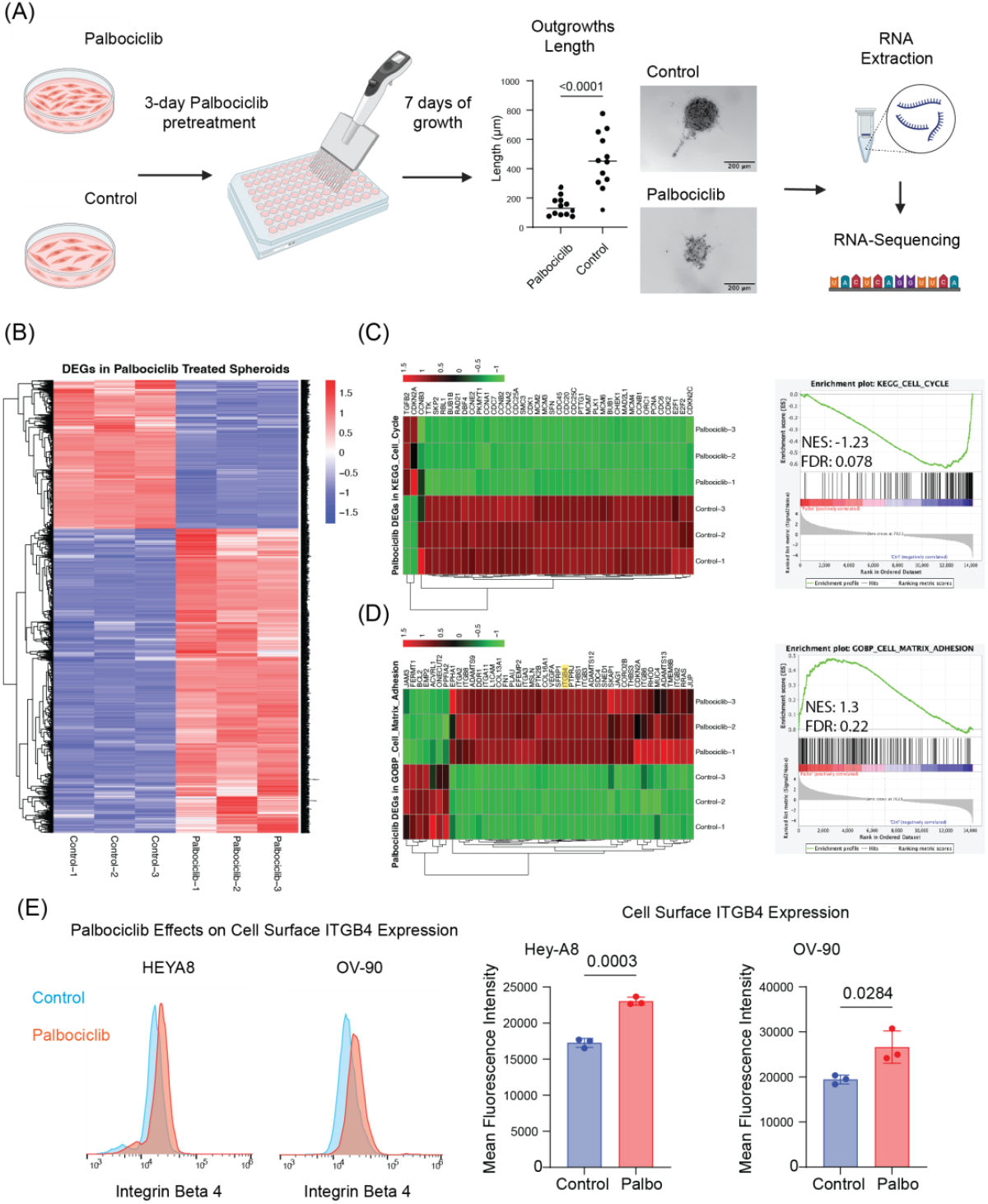
Palbociclib effects on ITGB4 expression and cell cycle genes in ovarian cancer 3D models. (**A**) Schematic of the experimental workflow where HEYA8 cells underwent pretreatment with 500 nM Palbociclib for 3 days, followed by 7 days of growth in 3D spheroid culture. The plot quantifies outgrowth length from the main spheroid in control versus Palbociclib-treated groups. Each point represents one spheroid, analyzed using an unpaired t-test. Representative images display spheroid formation in both treated and control groups, with subsequent steps including RNA extraction and sequencing. (**B**) Heatmap of differentially expressed genes (DEGs) comparing Palbociclib-treated spheroids to controls, highlighting over 2,600 DEGs. (**C**) Heatmap and GSEA plot demonstrating downregulation of cell cycle-related genes (KEGG_CELL_CYCLE pathway) in treated spheroids with a NES of -1.23 and an FDR of 0.078. (**D**) Heatmap and GSEA plot for the GOBP_CELL_MATRIX_ADHESION pathway showing an NES of 1.3 and an FDR of 0.224, illustrating differential expression patterns. (**E**) Flow cytometry and quantification of ITGB4 expression changes in HEYA8 and OV-90 cell lines post-Palbociclib treatment. Histograms show the differential cell surface ITGB4 levels, and bar charts quantify the changes in mean fluorescence intensity, with each dot representing an independent biological replicate. Statistical analysis was performed using an unpaired t-test, with p-values indicated on the charts.

Gene set enrichment analysis (GSEA) was performed based on RNA sequencing data to further investigate the effects of Palbociclib on gene expression in ovarian cancer cells which supports the illustrated heatmaps quantitatively. Especially, in (**Fig. 4C-right**), the GSEA plot versus KEGG_CELL_CYCLE pathway showed that cell cycle-related genes were significantly downregulated when treated with Palbociclib, documented with a normalized enrichment scores (NES) of -1.23 and a false discovery rate (FDR) q-value of 0.078. GOBP_CELL_MATRIX_ADHESION pathway analysis further provides an NES of 1.3 in (**Fig. 4D-right**), pointing toward the upregulation of genes involved in cell-ECM adhesion, with an FDR q-value of 0.224. These data highlight that Palbociclib treatment affects not only gene expression involved in the cell cycle but also significantly affects the gene expression involved in cell-matrix adhesion. We validated Palbociclib-regulation of integrin β4 expression in HEYA8 and OV90 cells using flow-cytometry-based quantification of cell surface protein expression (**Fig. 4E**).

Next, we wanted to determine whether Palbociclib-mediated suppression of cell proliferation and activation of cell-matrix adhesion programs is associated with reduced response to cisplatin. To do this we used a combination of flow cytometry and propidium iodide incorporation assay as a proxy for measuring cell death. In HEYA8 cells, which are characterized by low integrin β4 expression, Palbociclib significantly reduced the percentage of dead cells upon cisplatin treatment compared to the cisplatin treatment alone (p < 0.0001) (**Fig. 5A**). A similar trend was observed in OV-90 cells, where Palbociclib also significantly reduced cisplatin-induced cell death (p = 0.0116) (**Fig. 5B**). The decrease in cisplatin sensitivity was less marked in OV-90 cells, possibly because these cells express mid-level of integrin β4, therefore benefit less from integrin β4 upregulation compared to HEYA8 following Palbociclib pretreatment. The increased integrin β4 expression potentially provides a protective effect, enhancing the resistance to cisplatin-induced cell death. Furthermore, Gene Set Enrichment Analysis (GSEA) of RNA sequencing data from Palbociclib-treated spheroids demonstrated significant enrichment of cisplatin resistance gene sets across three independent studies (Kang^27^, Tsunoda^28^, and Whiteside^29^). The normalized enrichment scores (NES) for these studies were 1.35, 1.43, and 1.36, respectively, reinforcing the hypothesis that Palbociclib-induced growth arrest enhances cisplatin resistance (**Fig. 5C**).

**Figure 5.**
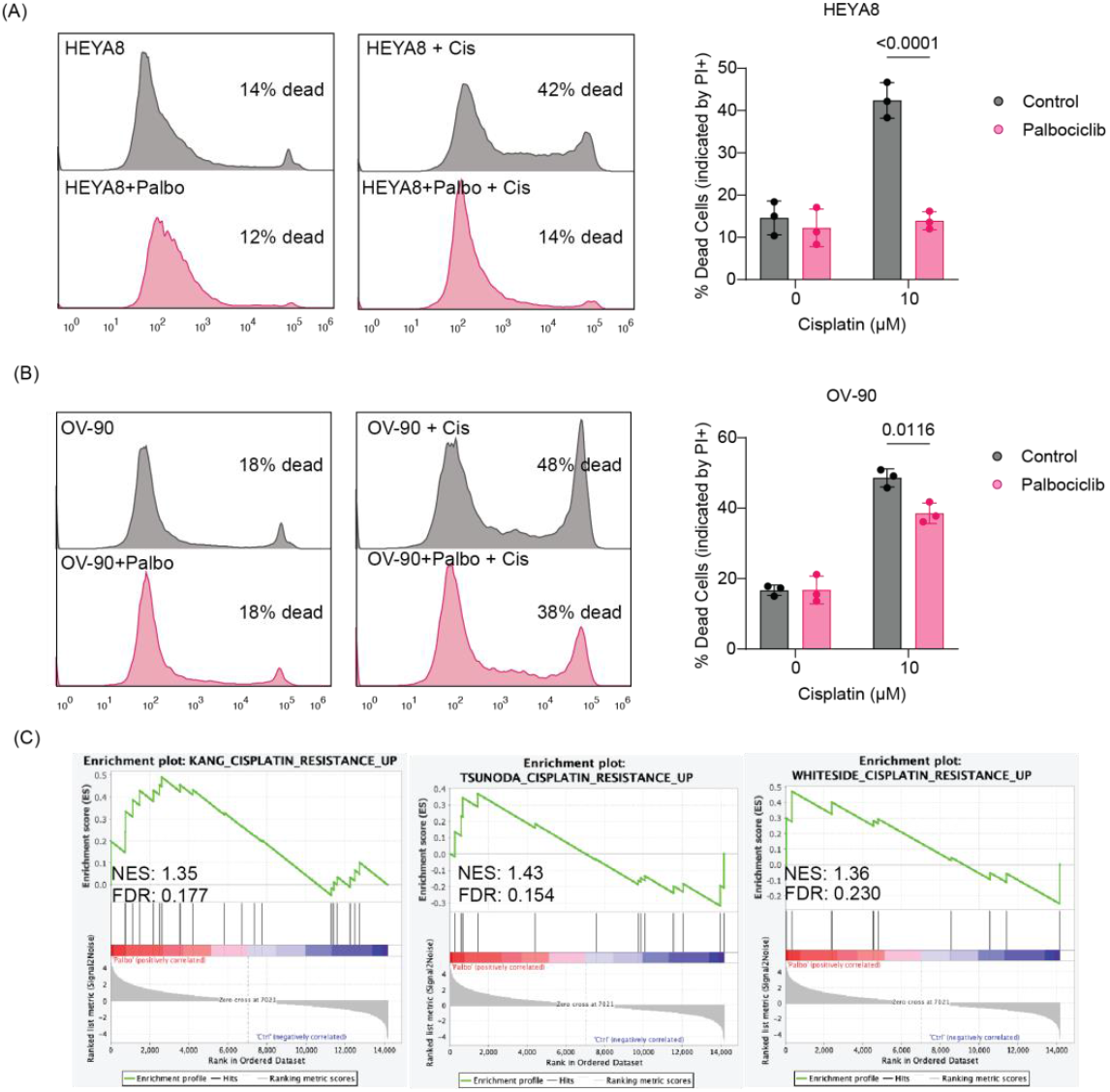
Palbociclib-induced cisplatin resistance in ovarian cancer cells. (**A**) Flow cytometry of HEYA8 cells shows reduced cell death when pretreated with Palbociclib (500 nM) before cisplatin (10 μM) treatment. Histograms show cell death percentages, with bar graphs quantifying the effects across treatments. Each dot represents an independent replicate. Statistical significance from decreased PI+ staining is shown (p < 0.0001), analyzed by two-way ANOVA highlighting interaction effects. (**B**) In OV-90 cells, similar pretreatment reduces cisplatin-induced cell death, with significant differences (p = 0.0116). Each bar graph dot represents a replicate, analyzed via two-way ANOVA. (**C**) GSEA of Palbociclib-treated HEYA8 spheroids shows upregulation of cisplatin resistance gene sets. NES and FDR for each study are Kang (NES: 1.35, FDR: 0.177), (NES: 1.43, FDR: 0.154), and Whiteside (NES: 1.36, FDR: 0.230), suggesting a link between Palbociclib pretreatment and altered drug response.

These findings suggest that Palbociclib enhances cisplatin resistance in ovarian cancer spheroids, with a particularly pronounced effect in low ITGB4-expressing cells, such as HEYA8, where the upregulation of integrin β4 may play a critical role in reducing cisplatin sensitivity.

### 3.5. Palbociclib Treatment Modulates Cisplatin Sensitivity in Patient-Derived Ovarian Cancer Organoids

We next sought to determine whether Palbociclib similarly induce a protective effect to cisplatin treatment in patient-derived ovarian cancer organoids (PDOs). Thus, we performed drug dose-response assays to cisplatin, comparing Palbociclib pre-treated vs. untreated PDOs. As expected, cisplatin treatment induced visible morphological changes in comparison to vehicle controls indicative of PDO sensitivity to cisplatin. Notably, these morphological changes were of lesser extent in Palbociclib pre-treated conditions (**Fig. 6A)**. To quantify responses to these treatments, we measured ATP levels as a surrogate of cell viability at the assay end-point using CellTiter-Glo® reagent revealing a robust, dose-dependent decrease in sensitivity to Cisplatin in Palbociclib-treated PDOs (**Fig. 6B**). These results suggest that the Palbociclib-induced resistance to cisplatin can be recapitulated in ovarian cancer patient primary material grown in the form of organoids.

**Figure 6.**
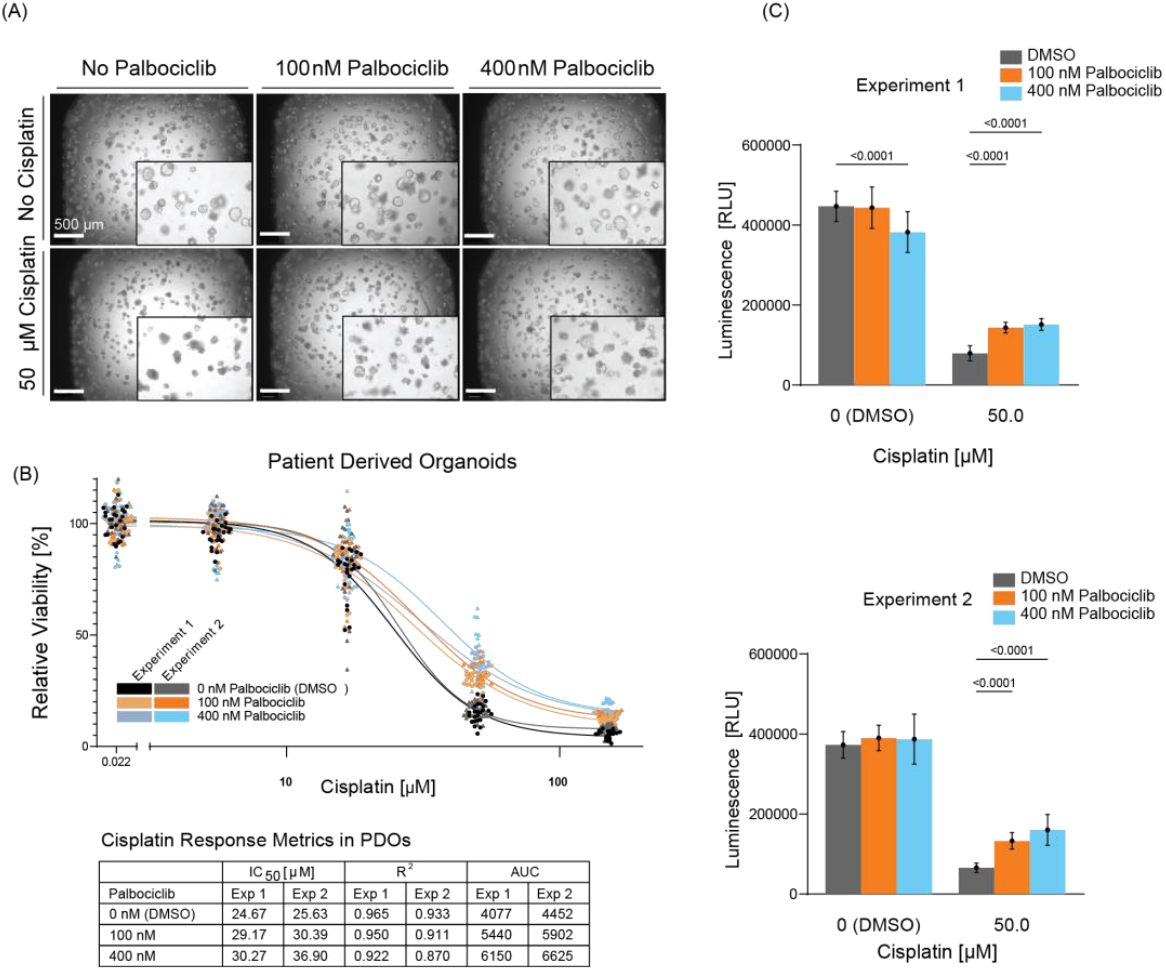
Palbociclib treatment decreases sensitivity to Cisplatin in patient-derived organoids (PDO). (**A**) Representative brightfield images at day 12 show control versus treated PDOs under different concentrations of Palbociclib and Cisplatin. Insets highlight morphological changes. Scale bars: 500 μm. (**B**) Drug dose-response curves of PDOs for cisplatin in untreated vs. palbociclib pre-treated PDOs at 100 nM and 400 nM concentrations. Each dot corresponds to one technical replicate (at least 9 replicates per dose in each independent experiment). Table shows IC50, R^2^, and AUC values for 50 μM cisplatin doses in two independent experiments. (**C**) Corresponding ATP levels (Relative Light Units, RLU) in PDOs treated with 50 μM Cisplatin after exposure to 100 nM or 400 nM Palbociclib. Levels were normalized to vehicle and Staurosporine-treated control (DMSO). Error bars represent mean ± SD of technical replicates. Statistical analysis utilized ordinary One-way ANOVA.

### 3.6. Overexpression of Integrin β4 Modulates Chemotherapy Response and Reduces Proliferation in Ovarian Cancer Cells

To determine whether integrin β4 expression modulates cisplatin sensitivity, we utilized a doxycycline-inducible lentiviral system to overexpress integrin β4 molecules in both HEYA8 and OV90 cells. As shown in **Fig. 7A**, we were able to overexpress ITGB4 in a doxycycline-dependent manner. Forced overexpression of integrin β4 significantly suppressed proliferation in both integrin β4-low HEYA8 and integrin β4-intermediate OV90 cells (p-value < 0.0001 and = 0.0008 respectively) albeit to a lesser extent in OV90 cells. These results suggest that the underlying basal levels of integrin β4 expression may modulate cell proliferation rates (**Fig. 7B**). We next measured the impact of integrin β4 overexpression on sensitivity to cisplatin using propidium iodide incorporation and flow cytometry. In both cell lines, integrin β4 overexpression exhibited significantly reduced Cisplatin-induced cell death compared to controls (**Fig. 7C-D**); hence, integrin β4 overexpression showed reduced drug sensitivity. Notably, these effects were of greater extent in HEYA8, the OC cell line with the lowest integrin β4 levels suggesting that the acquired expression of ITGB4 in low-expressing cells could have a significant impact in cisplatin resistance. These findings support the model whereby integrin β4 expression modulates ovarian cancer proliferation and response to chemotherapy (**Fig. 8**).

**Figure 7.**
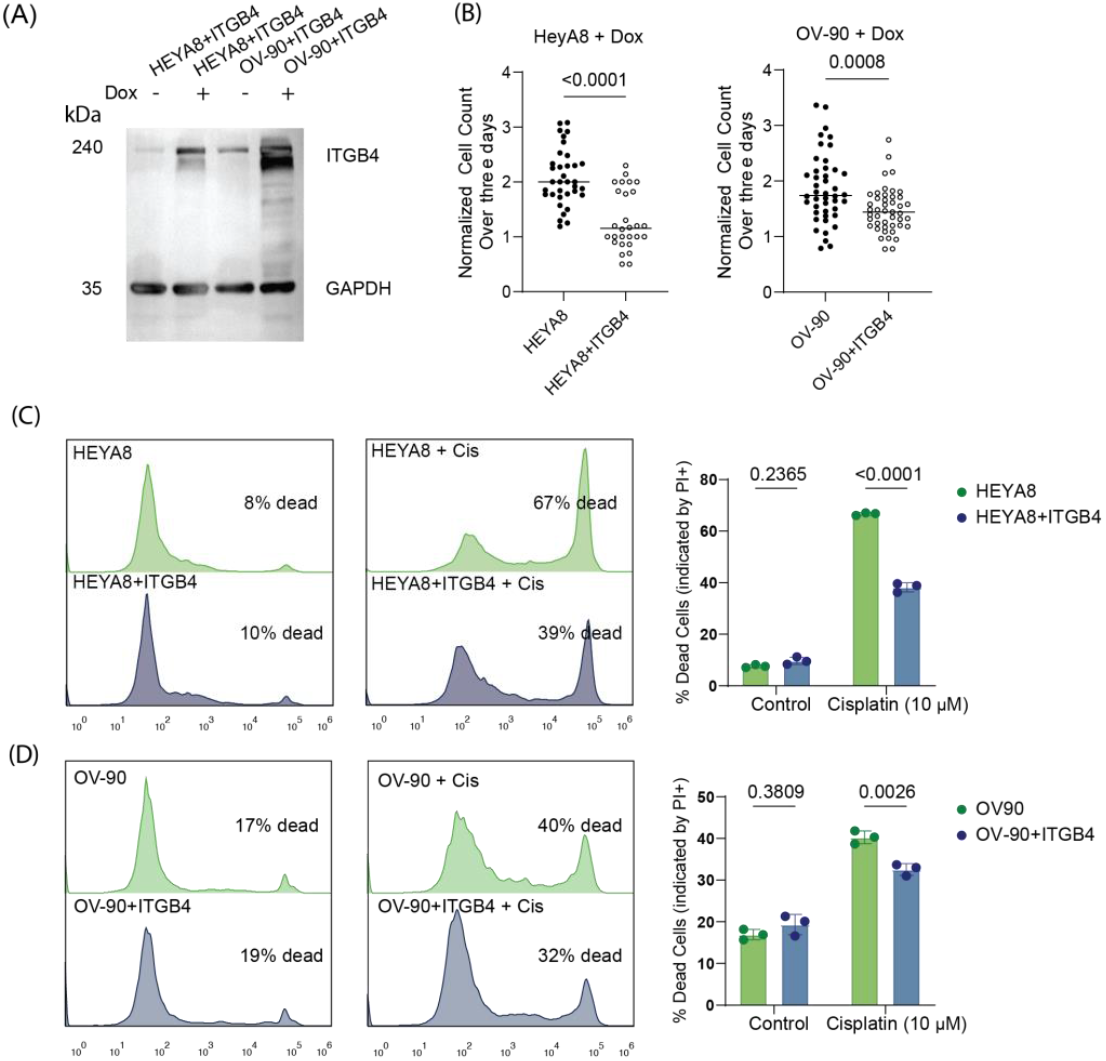
Integrin β4 modulates chemotherapy response and reduces proliferation in ovarian cancer cells. (**A**) Western blot analysis demonstrating induced overexpression of integrin β4 in HEYA8 and OV90 cells treated with doxycycline-inducible ITGB4 expression systems (doxITGB4). (**B**) Normalized cell count data showing significant suppression of proliferation due to induced overexpression of ITGB4 in HEYA8 and OV90 cells. Statistical significance determined by unpaired t-tests (p < 0.0001 for HEYA8, p = 0.0008 for OV90). (**C**) Flow cytometry analysis and bar graphs for HEYA8 cells show that ITGB4 overexpression reduces Cisplatin-induced cell death, with fewer dead cells in ITGB4-overexpressing cells compared to controls under Cisplatin treatment (p < 0.0001). Statistical analysis performed using two-way ANOVA test. (**D**) Similar analysis for OV90 cells, where ITGB4 overexpression confers increased resistance to Cisplatin. The comparison of native and ITGB4-overexpressing cells indicates a decrease in Cisplatin-induced cell death in ITGB4-overexpressing cells (p = 0.0026), with analysis supported by two-way ANOVA test and p-values are indicated on the chart.

**Figure 8.**
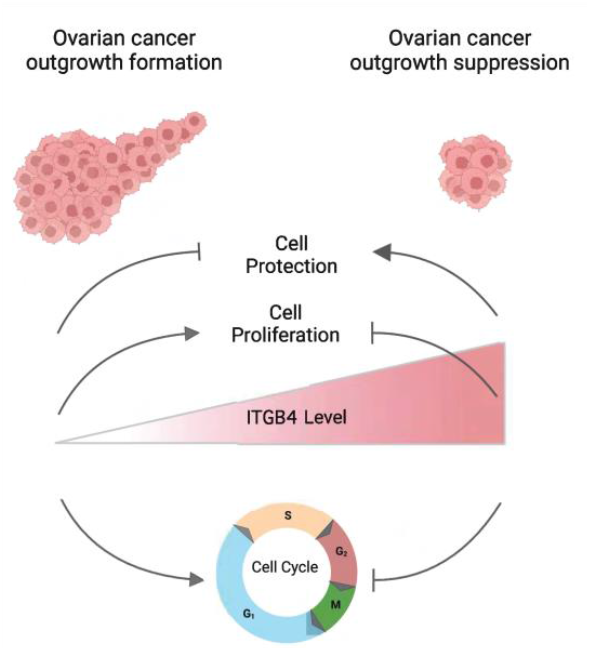
Influence of ITGB4 on Cell Cycle Dynamics and Cytotoxic Drug Resistance in Ovarian Cancer Models. This schematic illustrates the correlation between elevated ITGB4 expression and slowdown of cell cycle progression in various ovarian cancer models. Specifically, higher levels of ITGB4 are linked to reduced cellular proliferation, resulting in less outgrowth formation in 3D spheroid models and an increase in resistance to cytotoxic agents such as cisplatin.

## 4. Discussion

Our studies provide evidence that growth suppression of cultured OC cells decreases cisplatin sensitivity and induces integrin β4 expression. Furthermore, we demonstrated that forced expression of integrin β4 decreased proliferation and cisplatin sensitivity. Using patient data that are available through TCGA, we have demonstrated an inverse correlation between cell-matrix adhesion and cell-cycle progression. More specifically we identified that patient’s tumors expressing high levels of integrin β4 mRNA were the less proliferative, raising a question of whether regulation of integrin β4 expression is linked to cell proliferation. We provide experimental evidence that doxycycline-mediated regulation of integrin β4 expression can impact OC cell proliferation and their response to cisplatin, one of the most used chemotherapeutic agents in OC treatment.

Our in vitro experiments on OC cell lines were aligned with our analysis of patients’ data from TCGA; OC cell lines with higher integrin β4 protein levels displayed reduced proliferation rates, while those with lower expression proliferated more rapidly. However, it is important to note that some studies on other carcinomas have reported that higher expression levels of integrin α6β4 correlate with a greater proliferating fraction, suggesting context-dependent roles of integrin β4^30^ which could be related to the unique matrix compositions observed in different tumor contexts.

The relationship between growth rate of OC cells and cell-matrix adhesion were further elucidated by RNA-seq data of Palbociclib-treated OC spheroids. Treatment of OC cells with Palbociclib, a known inducer of G1 cell cycle arrest as a CDK4/6 inhibitor, resulted in the downregulation of cell cycle genes and an upregulation of genes involved in cell-matrix adhesion, interestingly including ITGB4 as one of the highly upregulated genes. Upregulation of Integrin β4 was confirmed by flow cytometry of Palbociclib-treated HEYA8 and OV-90 cells.

Palbociclib-treated cells had reduced sensitivity to the chemotherapeutic agent cisplatin. Pretreatment with Palbociclib profoundly diminished cisplatin-induced cell death in HEYA8 cells, which express low-levels of integrin β4, and less so in OV90 cells that expressed an intermediate level of integrin β4. These data support the idea that baseline integrin β4 expression modifies chemoresistance to cell cycle arrest induced by palbociclib treatment.

The efficacy of Palbociclib is closely associated with the functional status of the Rb protein because CDK4/6 inhibitors mediate the Rb pathway to exert a cell cycle arrest effect. Mutations or loss of Rb predispose cells to resistance against CDK4/6 inhibitors^31^. Indeed, previous studies have shown that cancer cell lines, lacking Rb expression, fail to respond to Palbociclib treatment^32 33 34^. HEYA8 and OV90 cells reportedly express functional Rb protein and thus should be sensitive to the inhibitory action of CDK4/6 ^35 36 37^. It is thus plausible that, following Palbociclib-mediated cell cycle arrest, the upregulation of integrin β4 can trigger survival signals which diminish the sensitivity of the cells to cisplatin cytotoxicity. These findings comply with reports that integrin β4 promotes cell survival and resistance to apoptotic-inducing agents^8 7^.

The interaction between upregulated integrin β4 and laminin in the ECM may increase survival signals and activate intracellular signaling pathways that promote cell survival and inhibit apoptosis ^5 6^. The α6β4 heterodimer interacts with laminin and can activate signaling cascades such as PI3K/Akt and NF-κB pathways^38 39^. These pathways are associated with cell survival, invasion, and chemotherapy resistance. Based on this, we hypothesize that the cell cycle is inversely correlated with Cell-ECM adhesion including integrin β4, and the higher expression of this adhesion molecule suppresses sensitivity to cisplatin. These findings also suggest that cell cycle inhibition can induce a phenotypic shift towards a more quiescent phenotype, with increased cell adhesion mediated by integrin β4^40^. These cells are more resistant to treatments that target rapidly dividing cells^41^. We propose a model in which elevated integrin β4 expression slows cell cycle progression and reduces proliferation, thereby increasing resistance to cytotoxic agents like cisplatin (**Fig. 8**).

In our in vitro study with OC cell lines, high levels of integrin β4 led to lower proliferation and increased cisplatin resistance. Additionally, integrin β4 has been shown to protect breast cancer cells from DNA damage-induced apoptosis via activation of survival pathways ^8 39^.

Based on our results, integrin β4 expression in tumors may act as a biomarker for patient stratification at risk for poor chemotherapy outcomes, and personalized treatment strategies would be warranted. Also, targeting pathways mediated by integrin β4 can enhance chemosensitivity, yielding better results in combination with standard therapeutic agents like Palbociclib^32 42^. Further studies are needed to fully elucidate the molecular mechanisms whereby chemoresistance is conferred by integrin β4 and to identify intervention targets^43^. In vivo studies will be essential for establishing the rationale for targeting integrin β4 in ovarian cancer. Similarly, our study acknowledges limitations regarding the use of cell lines and organoids, which cannot fully reflect the complexity of the tumor microenvironment.

## 5. Conclusions

In summary, our study suggests that integrin β4 is an important modulator of proliferation and response to cisplatin in ovarian cancer cell models. Expression of integrin β4 inversely correlates with cell cycle progression; the ability of integrin β4 to enhance the survival of cells undergoing cell cycle arrest contributes to chemoresistance and points to a potential target for improving patient outcomes.

## Author Contributions

S.F.: Conceptualization, Methodology, Validation, Software, Formal analysis, Investigation, Data curation, Writing—original draft, Visualization. D.C.: Methodology, Validation, Formal analysis, Investigation, Visualization. J.-S.C.: Conceptualization, Methodology, Formal analysis, Investigation, Writing. T.P. (Teagan Polotaye): Methodology, Validation, Formal analysis, Investigation, Visualization. T.P. (Tonja Pavlovic): Conceptualization, Methodology. P.B.: Methodology, Software, Formal Analysis, Visualization. T.M.: Conceptualization, Methodology, Validation. P.L.: Conceptualization, Software. L.A.M.: Conceptualization, Methodology, Funding acquisition. M.P.I.: Conceptualization, Methodology, Writing—original draft, Visualization, Supervision, Funding acquisition. All authors have read and agreed to the published version of the manuscript.

